## Supplementary Information for "A way to break bones? The weight of intuitiveness"

Supporting Informations

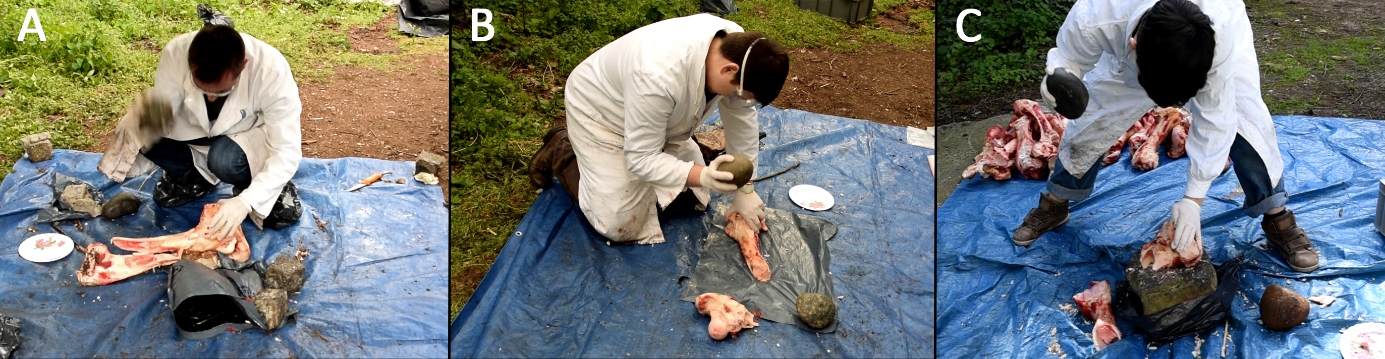

Supporting Information 1: Example of individual position during the breakage. A-individual n°10 squatting, B-individual 8 kneeling and C-individual n°1 standing.

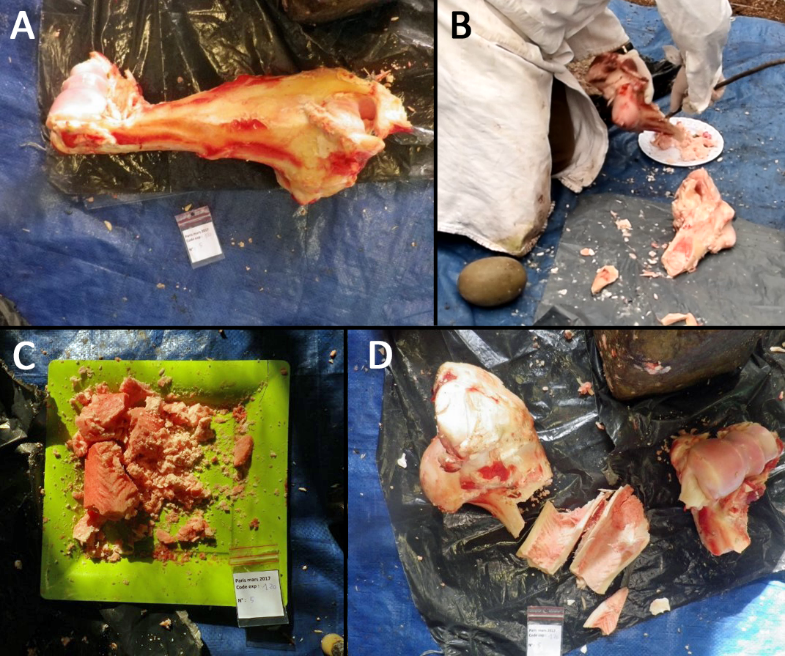

Supporting Information 2: Bone number 5 of individual n°7 (A, C and D) and Individual 8 (B); A-before breakage, B-yellow marrow recovery with wood stick, C-marrow recovered and D-bone remains after breakage.

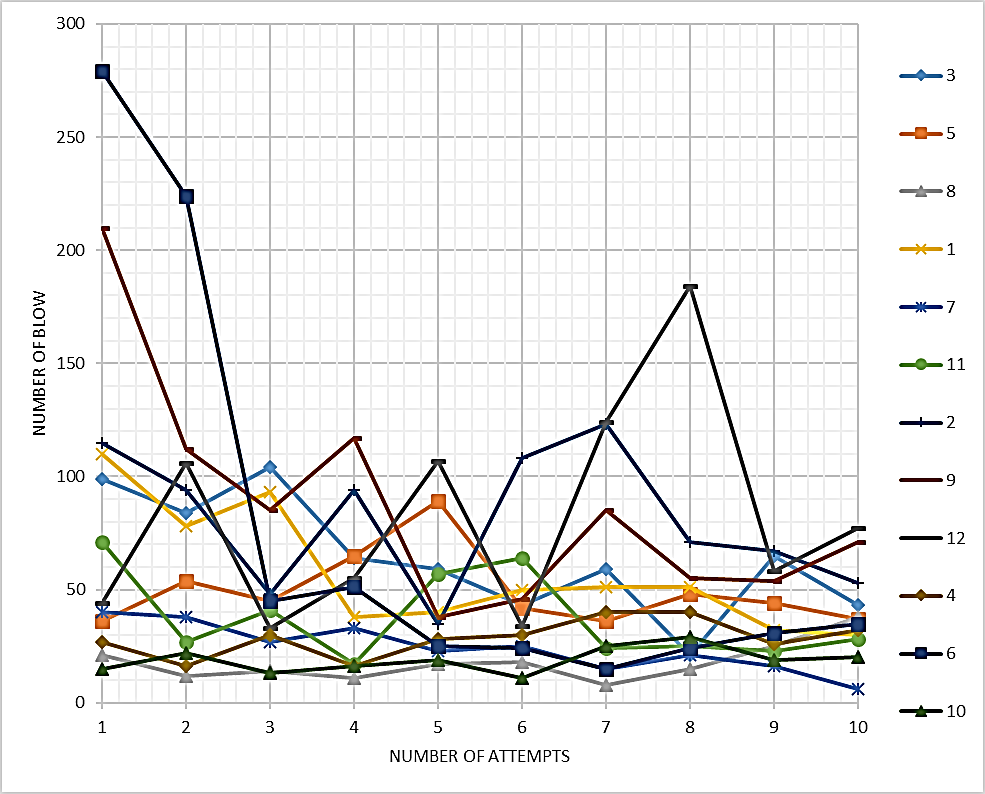

Supporting Information 3: The number of blows according to the number of try for each individual.

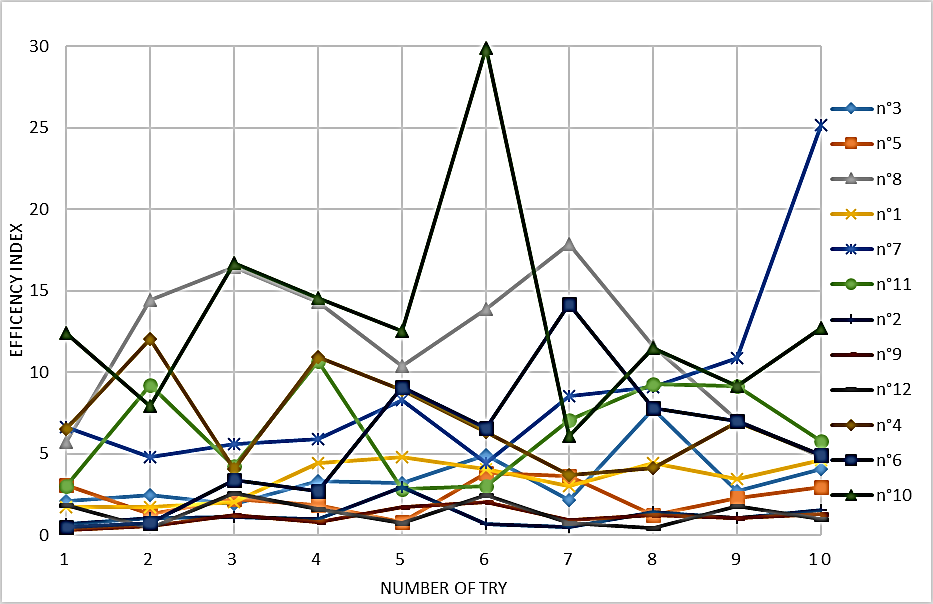

Supporting Information 4: Efficiency Index by individual during the experiment for each bone.

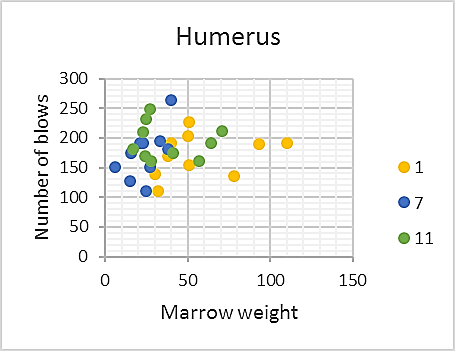

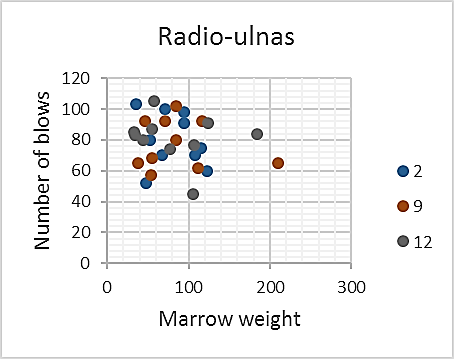

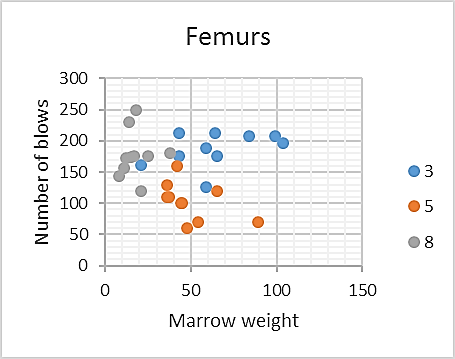

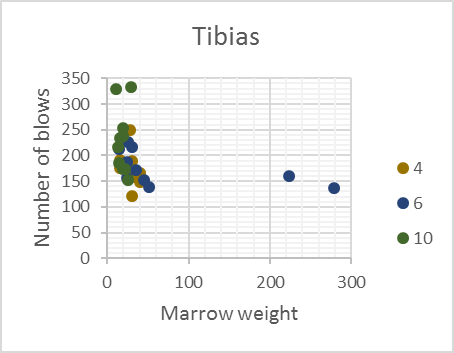

Supporting Information 5: Graph of the number of blows by the marrow weight (in gram) for each element

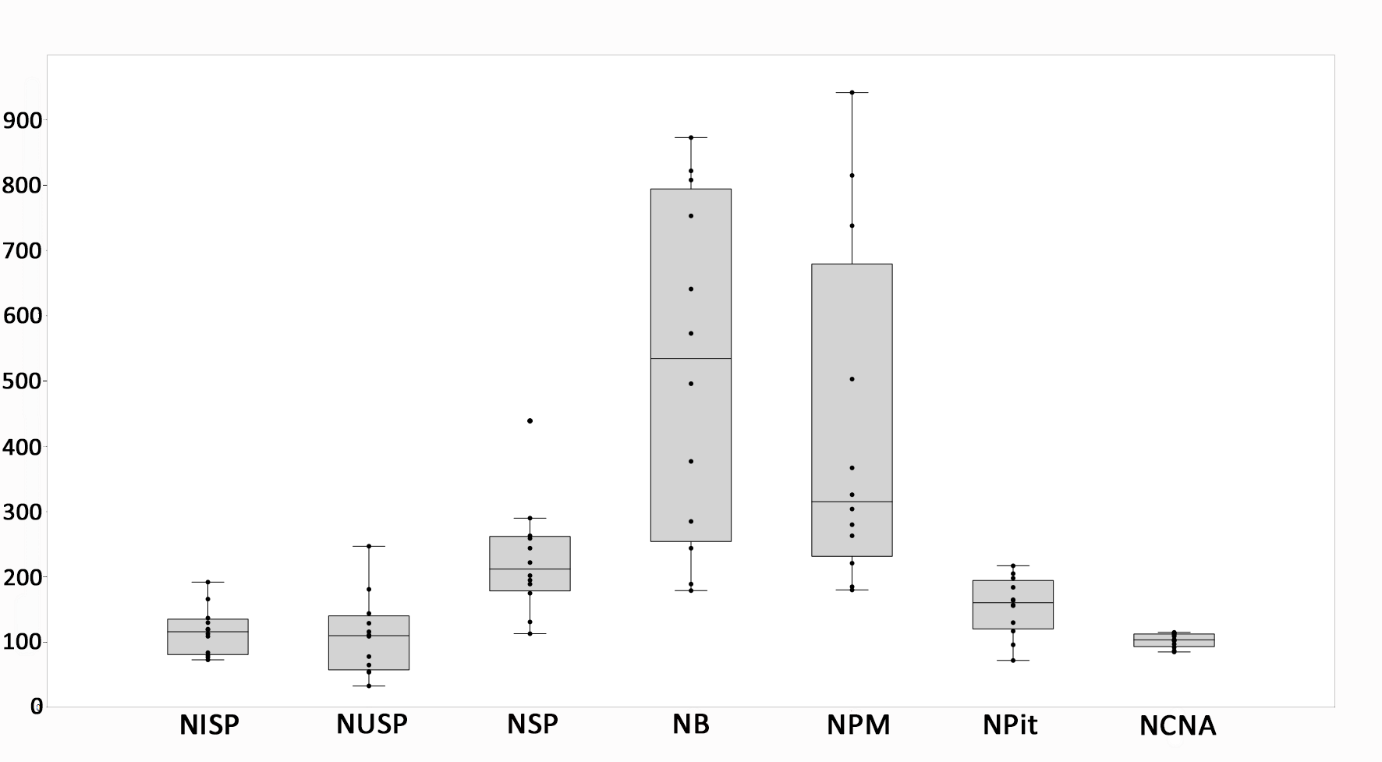

Supporting Information 6: Box plots and jitter with outsider of marrow Weight (MW), Number of Specimen (NSP), Number of identified specimens (NISP), Number of undetermined specimens (NUSP), Number of blows (NB), Number of percussion marks (NPM), Number of pit and grooves (Npit) and Number of crushing marks, of adhering flakes and of notches (NCNA).

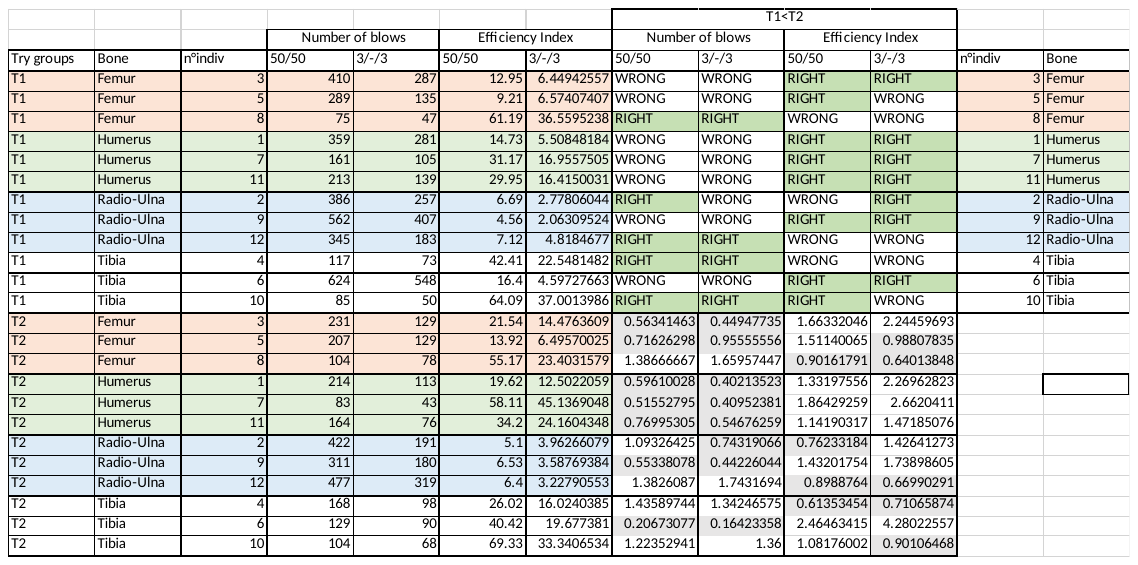

Supporting Information 7: The number of blows and the Efficiency Index, for each individual, regarding 50/50 the five first tries (T1) and the five last ones (T2), 3/-/3 the three first tries (T1) and the three last ones (T2).

|  |  | All Element | Humerus | Radio-ulna | Tibia | Femur |
| --- | --- | --- | --- | --- | --- | --- |
| Number of blows (50/50) | Degree of freedom | 23 | 2 | 2 | 2 | 2 |
|  | t-test assumption | No | No | No | No | No |
|  | Wilcoxon signed rank test | 50 | 6 | 3 | 3 | 5 |
|  | P-value | 0.1294 | 0.25 | 1 | 1 | 0.5 |
| Number of blows  (3/-/3) | Degree of freedom | 23 | 2 | 2 | 2 | 2 |
|  | t-test assumption | No | No | No | No | No |
|  | Wilcoxon signed rank test | 61 | 6 | 4 | 3 | 4 |
|  | P-value | 0.09229 | 0.0254 | 0.75 | 1 | 0.75 |
| Efficiency Index (50/50) | Degree of freedom | 23 | 2 | 2 | 2 | 2 |
|  | t-test assumption | No | No | No | No | No |
|  | Wilcoxon signed rank test | 21 | 0 | 3 | 2 | 2 |
|  | P-value | 0.1763 | 0.25 | 1 | 0.75 | 0.75 |
| Efficiency Index  (3/-/3) | Degree of freedom | 23 | 2 | 2 | 2 | 2 |
|  | t-test assumption | No | No | No | No | No |
|  | Wilcoxon signed rank test | 26 | 0 | 3 | 3 | 4 |
|  | P-value | 0.3394 | 0.25 | 1 | 1 | 0.75 |

Supporting Information 8: Results of Wilcoxon signed rank test of the Number of blows and the Efficiency Index between the first five attempts and the last five ones (50/50) and between the first three attempts and the last three ones (3/-/3).

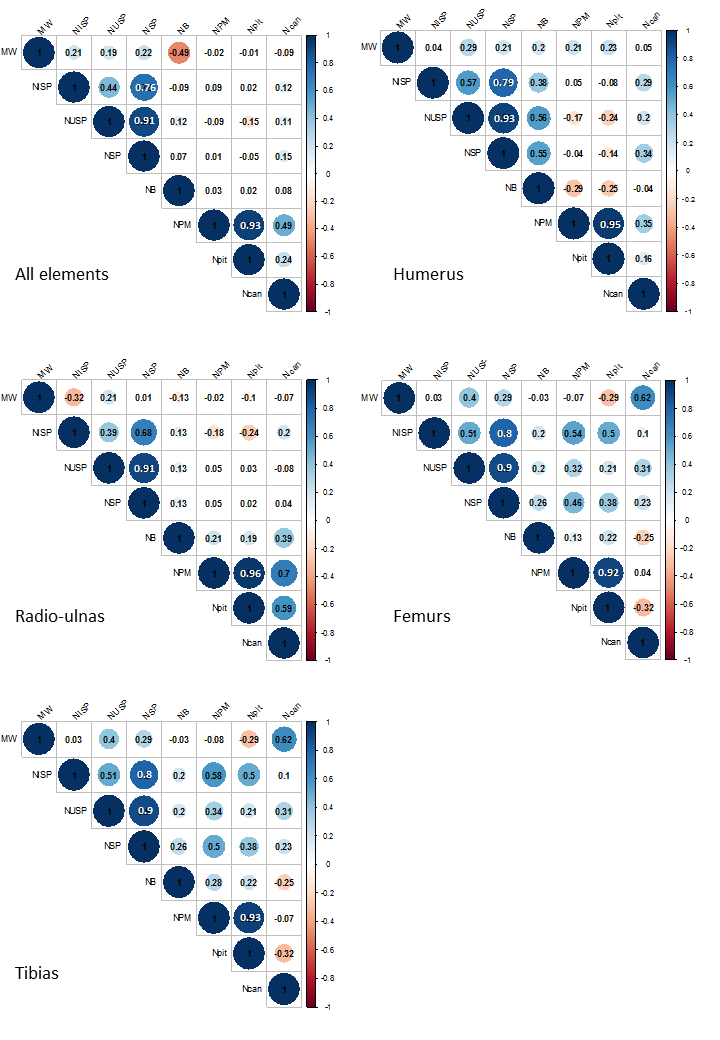

Supporting Information 9: Spearman's rank correlation coefficient results between the extracted marrow Weight (MW), Number of Specimen (NSP), Number of identified specimens (NISP), Number of undetermined specimens (NUSP), Number of blows (NB), Number of percussion marks (NPM), Number of pit and grooves (Npit) and Number of crushing marks, of adhering flakes and of notches (NCNA) for all elements, and each elements separately.

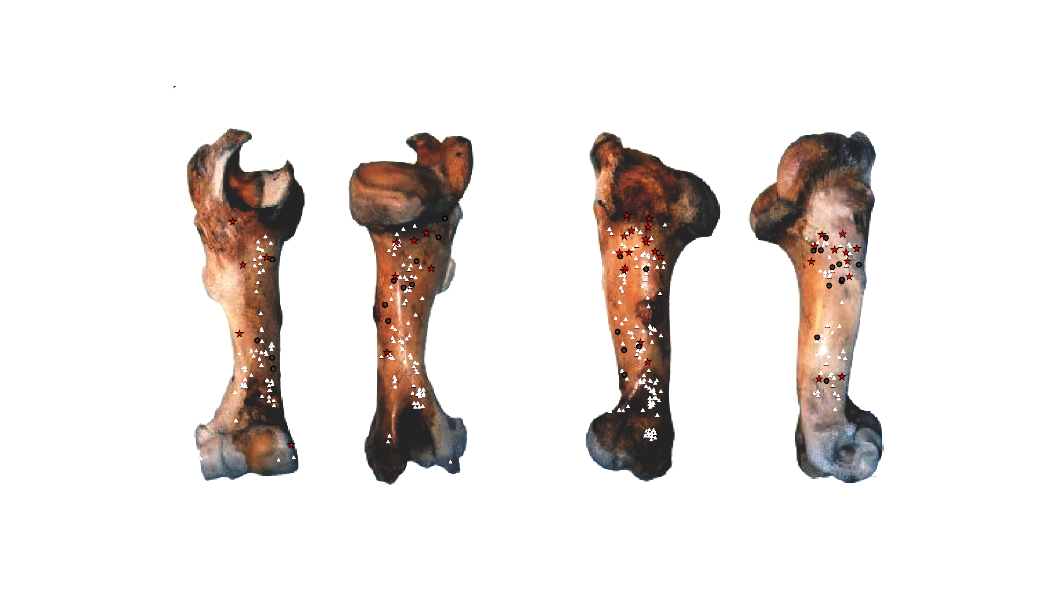

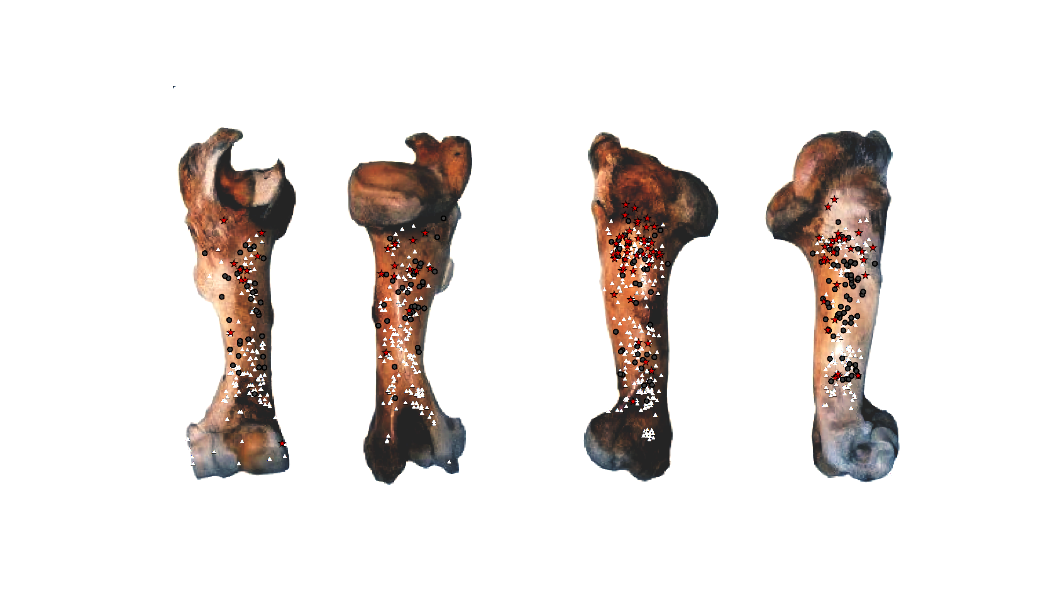

C

Ant. Post.	Med.	Lat.

Ant. Post.	Med.	Lat.

D

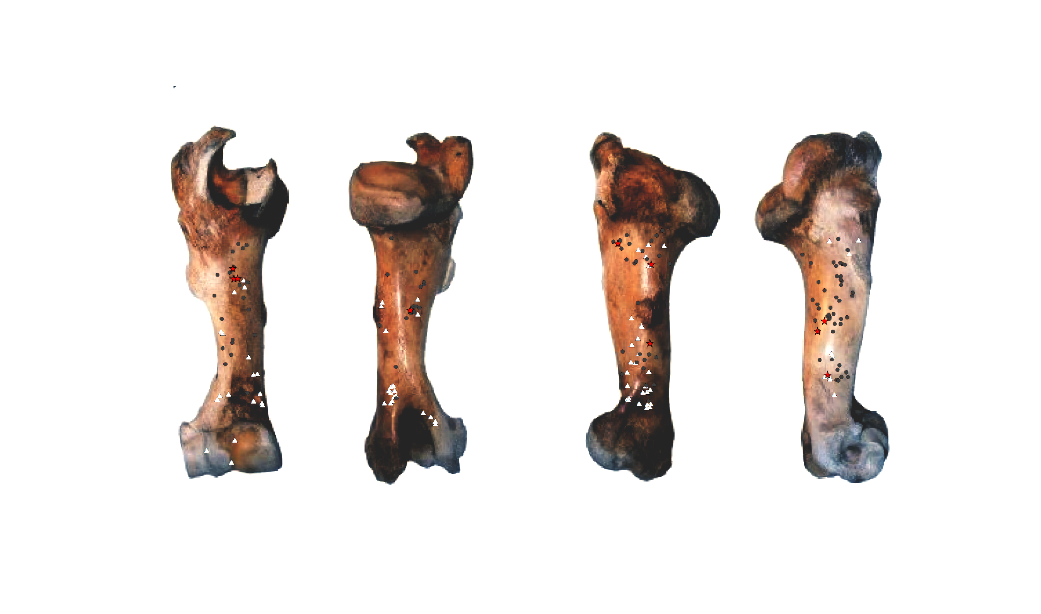

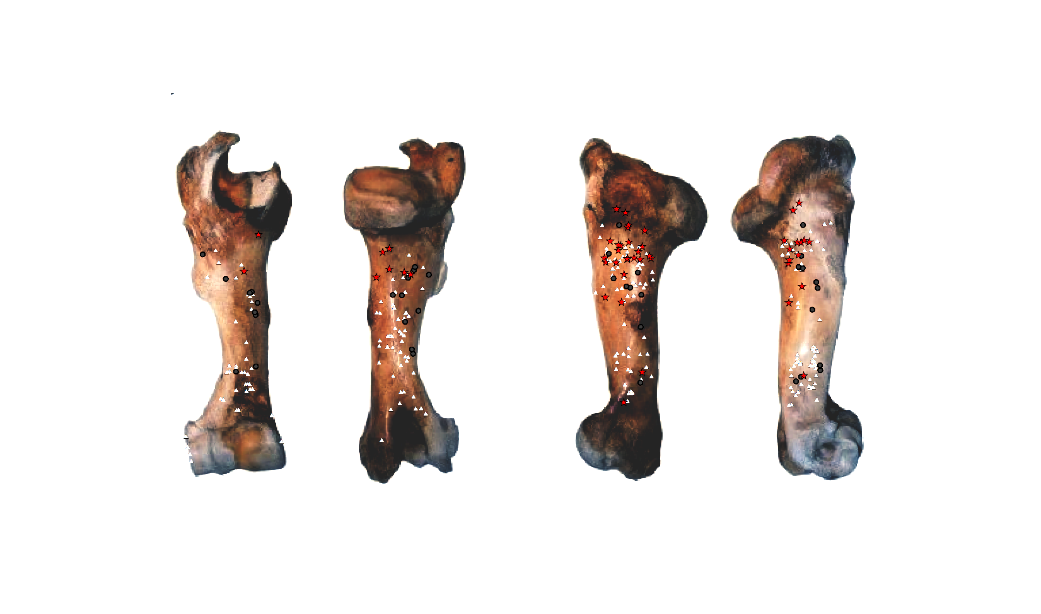

Ant. Post.	Med.	Lat.

Ant. Post.	Med.	Lat.

A

B

Supporting Information 10: Distribution of percussion marks along the humerus divided by type of percussion mark (red star: crushing marks, white triangle: pits and grooves and grey circle: notches and adhering flakes); A- Individual n°1; B- Individual n°7; C- Individual n°11; D- Merged humerus series.

Supporting Information 11: Distribution of percussion marks along the radio-ulnas divided by type of percussion mark (red star: crushing marks, white triangle: pits and grooves and grey circle: notches and adhering flakes); A- Individual n°2; B- Individual n°9; C- Individual n°12; D- Merged radio-ulnas series.

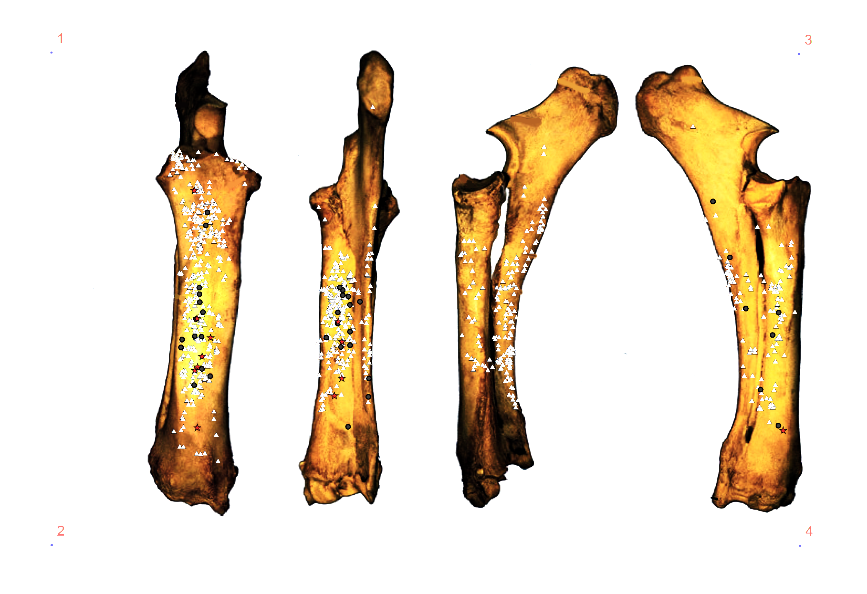

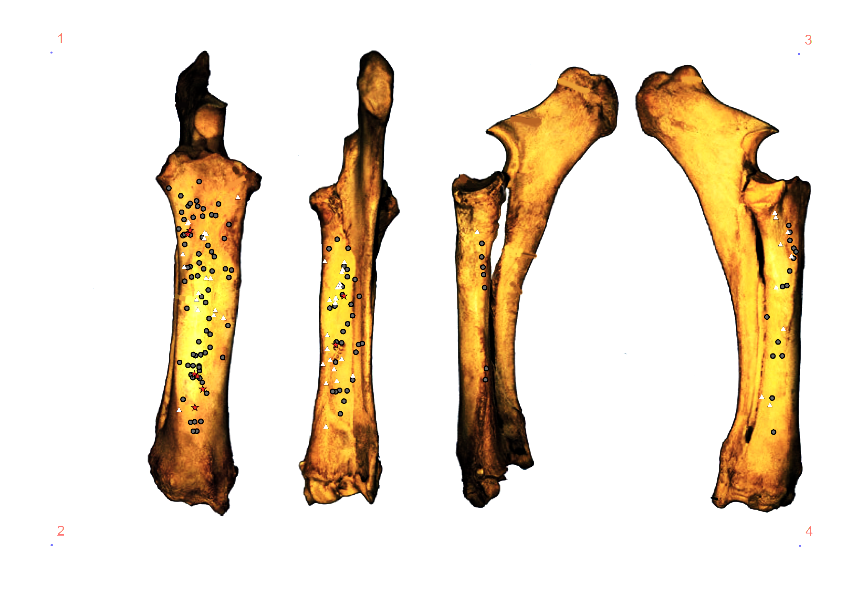

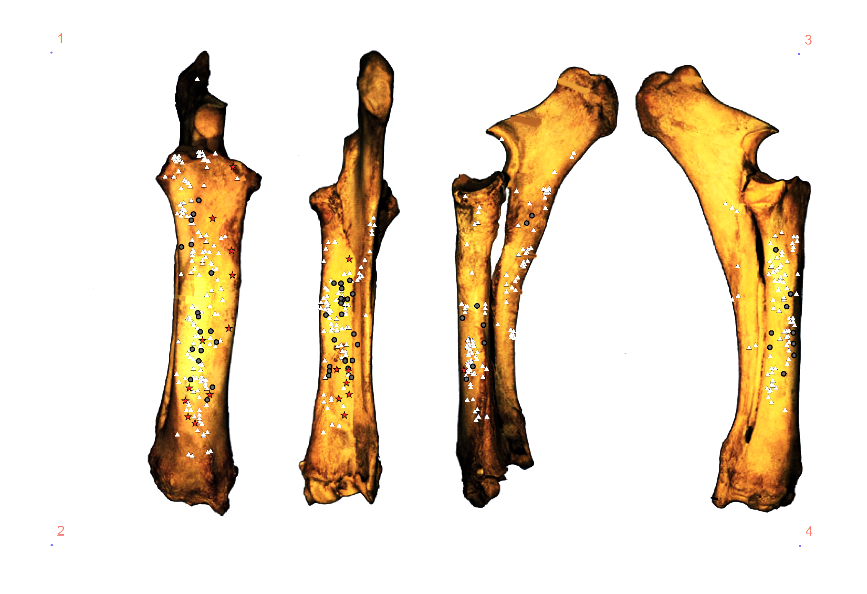

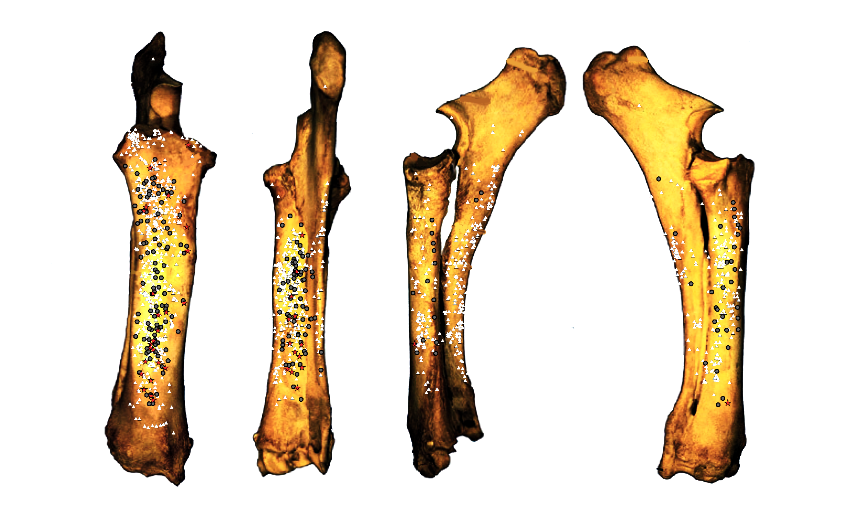

A

B

C

D

Ant. Post. Med.	Lat.

Ant. Post	Med.	Lat.

Ant. Post. Med.	Lat.

Ant. Post. Med.	Lat.

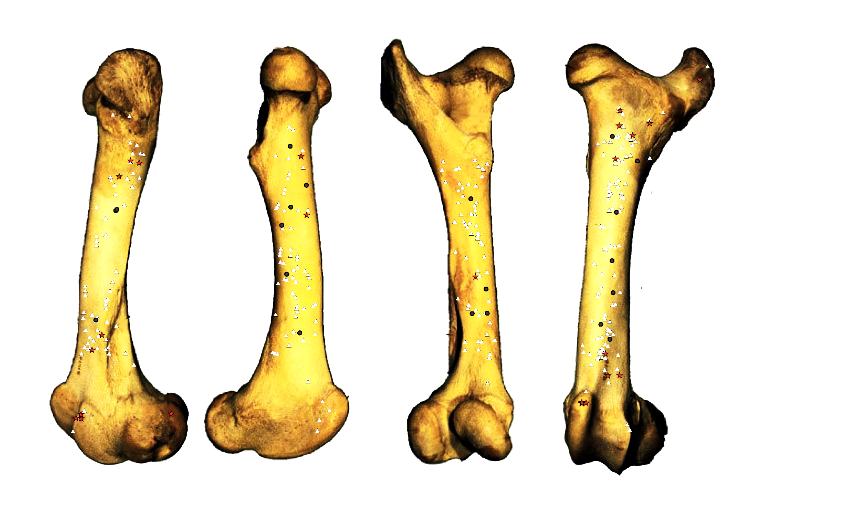

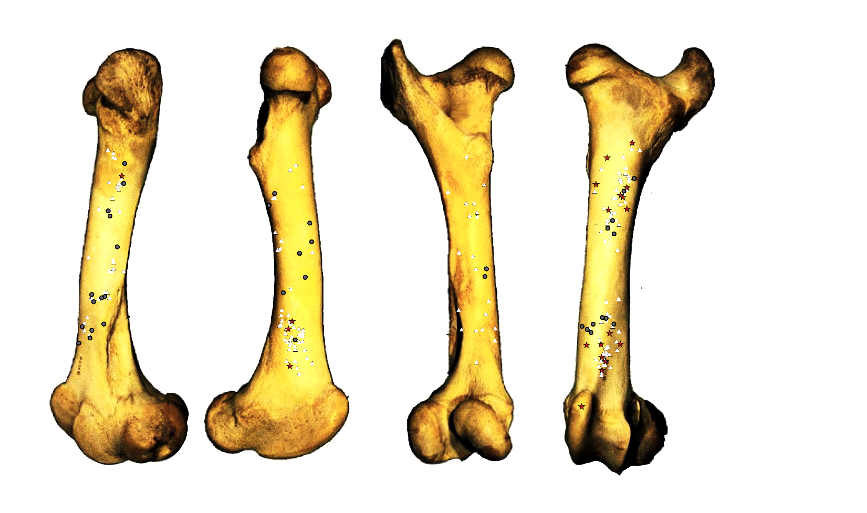

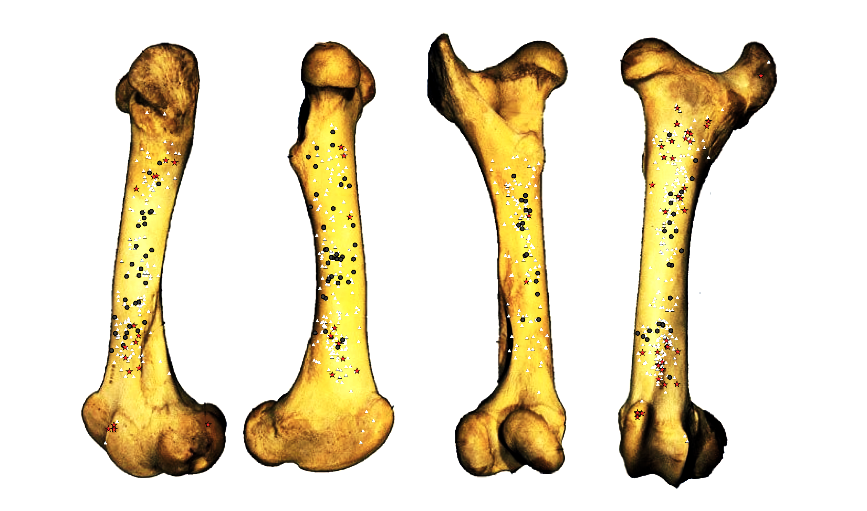

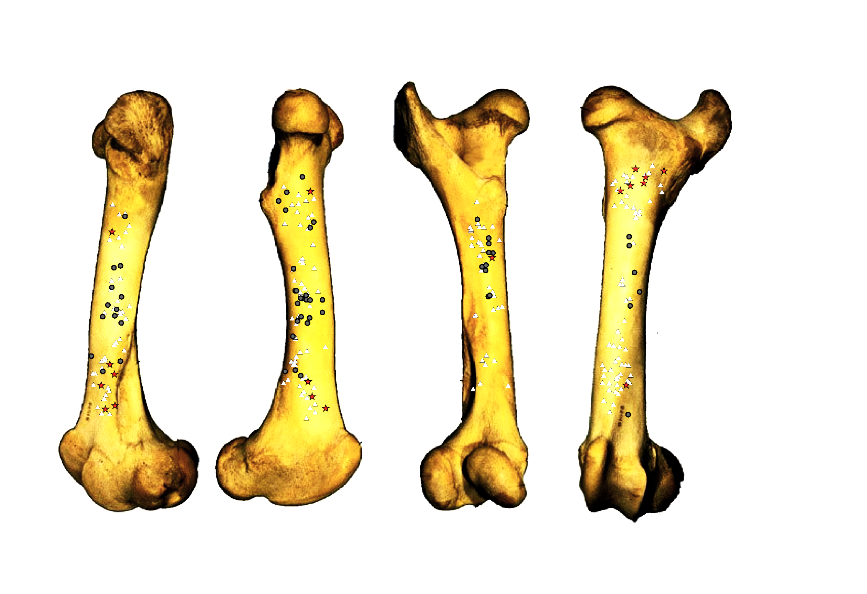

A

B

C

D

Ant. Post.	Med. Lat.

Ant. Post. Med.	Lat.

Ant. Post. Med.	Lat.

Ant. Post.	Med. Lat.

Supporting Information 12: Distribution of percussion marks along the femurs divided by type of percussion mark (red star: crushing marks, white triangle: pits and grooves and grey circle: notches and adhering flakes); A- Individual n°3; B- Individual n°5; C- Individual n°7; D-: Merged femurs.

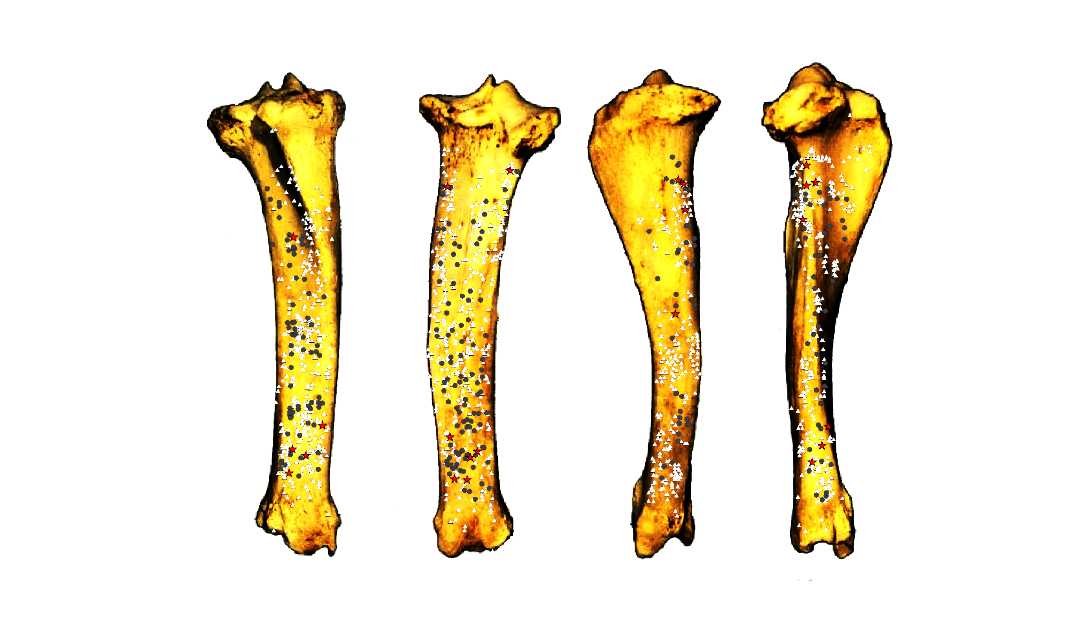

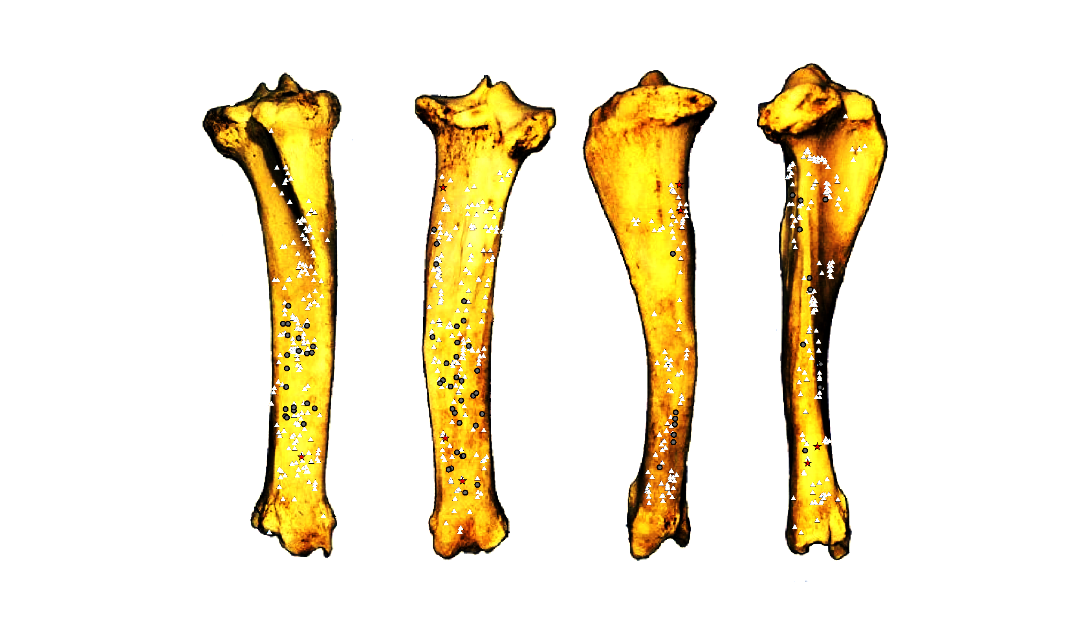

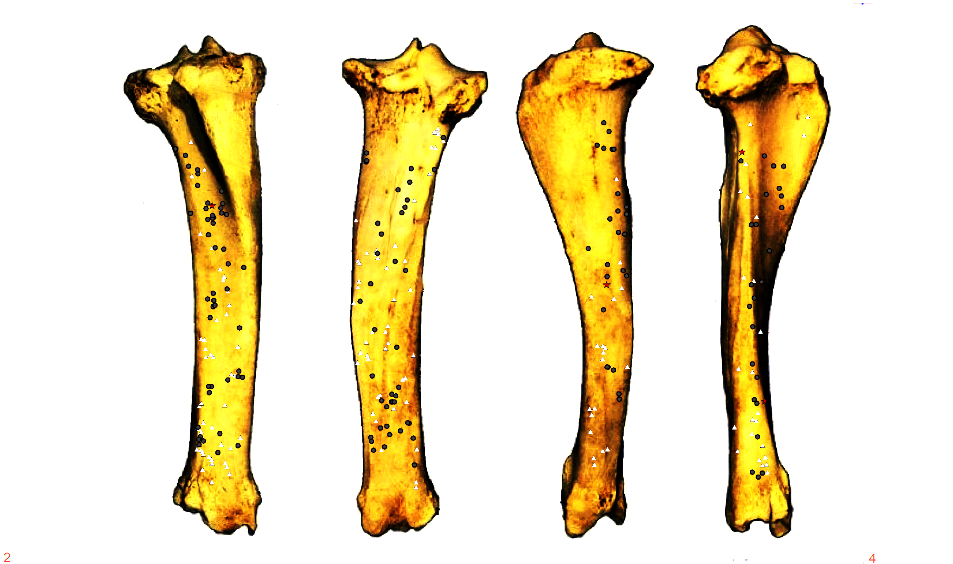

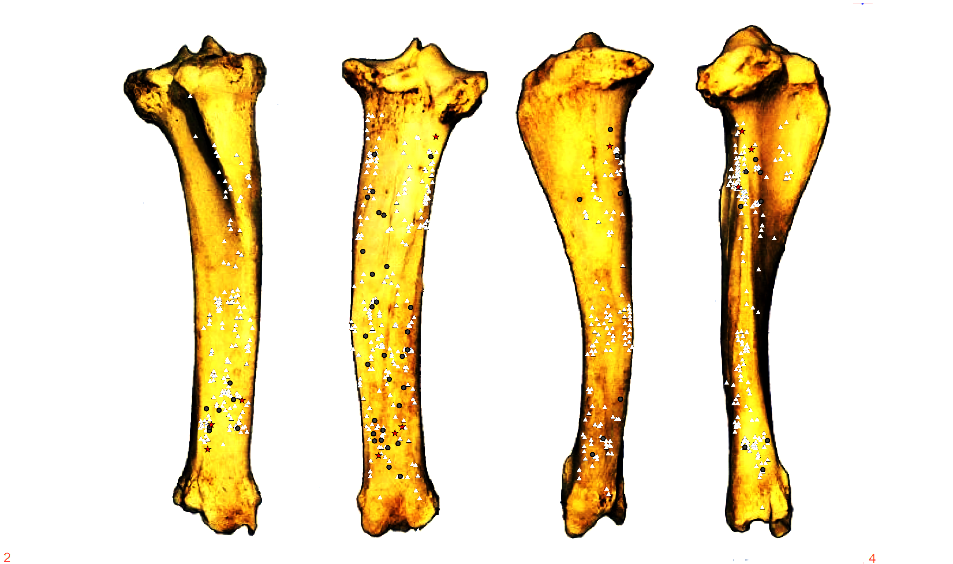

A

B

C

D

Ant. Post.	Med. Lat.

Ant. Post. Med.	Lat.

Ant. Post. Med.	Lat.

Ant. Post.	Med. Lat.

Supporting Information 13: Distribution of percussion marks along tibias divided by type of percussion mark (red star: crushing marks, white triangle: pits and grooves and grey circle: notches and adhering flakes A- Individual n°4; B- Individual n°6; C- Individual n°10; D- Merged tibias series.

Supporting Information 14: Vettese, Delphine; Stavrova, Trajanka; Borel, Antony; Marín, Juan; Moncel, Marie-Hélène; Arzarello, Marta; et al. (2020): SI 14: Dataset of observations during the intuitive experiment of long bone breakage. figshare. Dataset. <https://doi.org/10.6084/m9.figshare.12709592>

Supporting Information 15: Vettese, Delphine; Stavrova, Trajanka (2020): SI 15: Zooarchaeological dataset of observations during the analyses of long bone remains coming from the intuitive experiment of long bone breakage. figshare. Dataset. <https://doi.org/10.6084/m9.figshare.12709709.v1>

Supporting Information 16: Experiment protocol: Delphine Vettese 2020. An archaeological experiment focused on the intuitive way of long bones breakage to extract marrow. **protocols.io** [dx.doi.org/10.17504/protocols.io.be7fjhjn](https://www.protocols.io/view/an-archaeological-experiment-focused-on-the-intuit-be7fjhjn/dx.doi.org/10.17504/protocols.io.be7fjhjn)
